# PERK Activation by SB202190 Attenuates Amyloidogenesis via TFEB-induced Autophagy-Lysosomal Pathway

**DOI:** 10.1101/2021.05.25.445568

**Authors:** Mihyang Do, Jeongmin Park, Yubing Chen, So-Young Rah, Thu-Hang Thi Nghiem, Jeong Heon Gong, Byung-Sam Kim, Jeong Woo Park, Stefan W. Ryter, Young-Joon Surh, Uh-Hyun Kim, Yeonsoo Joe, Hun Taeg Chung

## Abstract

The protein kinase R (PKR)-like endoplasmic reticulum (ER) kinase (PERK), a key ER stress sensor of the unfolded protein response (UPR), can confer beneficial effects by facilitating the removal of cytosolic aggregates through the autophagy-lysosome pathway (ALP). In neurodegenerative diseases, the ALP ameliorates the accumulation of intracellular protein aggregates in the brain. Transcription factor-EB (TFEB), a master regulator of the ALP, positively regulates key genes involved in the cellular degradative pathway. However, in neurons, the role of PERK activation in mitigating amyloidogenesis by ALP remains unclear. In this study, we found that SB202190 selectively activates PERK independently of its inhibition of p38 mitogen-activated protein kinase, but not inositol-requiring transmembrane kinase/endoribonuclease-1α (IRE1α) or activating transcription factor 6 (ATF6), in human neuroblastoma cells. PERK activation by SB202190 was dependent on mitochondrial ROS production and promoted Ca^2+^-calcineurin activation. The activation of the PERK-Ca^2+^-calcineurin axis by SB202190 positively affects TFEB activity to increase ALP in neuroblastoma cells. Collectively, our study reveals a novel physiological mechanism underlying ALP activation, dependent on PERK activation, for ameliorating amyloidogenesis in neurodegenerative diseases.

## Introduction

Amyloids are a highly organized form of protein aggregation typically associated with human neuropathies, including Alzheimer’s diseases (AD), Parkinson’s diseases (PD), and Huntington’s diseases (HD) (Knowles *et al*, 2014). Under various pathological conditions, amyloid fibrils arise from amyloidogenesis as a biochemical process in which a soluble protein is converted into insoluble and fibrillar protein aggregates (Chiti & Dobson, 2006; Sipe, 1992). Accumulated aggregated proteins, such as amyloid β (Aβ), can trigger endoplasmic reticulum (ER) stress and lead to many neurodegenerative diseases (Bell *et al*, 2016; Hughes & Mallucci, 2019; Ron & Walter, 2007; Rozpedek *et al*, 2015). ER stress occurring in neurodegenerative diseases activates all three pathways of the mammalian unfolded protein response (UPR), each represented by its unique UPR sensor: PKR-like ER kinase (PERK) (Harding *et al*, 1999), inositol-requiring transmembrane kinase/endoribonuclease-1α (IRE1α) (Cox *et al*, 1993) and activating transcription factor 6 (ATF6) (Wang *et al*, 2000). However, it has become evident that the PERK pathway exerts a major role in the resolution of aggregated protein cytotoxicity. PERK activation by ER stress mitigates cerebral Aβ accumulation by reducing β-secretase-1 levels and cognition deficits in AD (Devi & Ohno, 2014; Ma *et al*, 2013). Moreover, UPR induction leads to the accumulation α-synuclein (α-syn) that contributes to neurodegeneration in α-synucleinopathies (Colla *et al*, 2012). Beyond its canonical role in the UPR, PERK as a structural tether of the mitochondria-associated ER membrane (MAMs), serves as a physical and functional connection between the ER and the mitochondria (Hayashi *et al*, 2009; Verfaillie *et al*, 2012). PERK signaling facilitates Ca^2+^ efflux from the ER, and increases mitochondrial Ca^2+^ levels, leading to increased generation of mtROS (Zhang *et al*, 2019). The increase of cytosolic Ca^2+^ levels by PERK (Zhang *et al*., 2019) activates calcineurin, a Ca^2+^ and calmodulin-dependent phosphatase (Bollo *et al*, 2010). In particular, the interaction of PERK and calcineurin in astrocytes promotes cell survival after acute brain injuries (Chen *et al*, 2016).

Neurodegenerative disease is strongly associated with defects in the autophagy-lysosomal pathway (ALP) (Menzies *et al*, 2015). Moreover, the clearance of intracellular aggregates in the brain typically improves symptoms of neurodegenerative diseases such as AD, PD, and HD (Martini-Stoica *et al*, 2016; Yamamoto & Simonsen, 2011). ALP has been widely demonstrated to ameliorate pathology in these diseases (Decressac *et al*, 2013; Tsunemi *et al*, 2012; Xiao *et al*, 2015). TFEB, a global regulator of ALP, is a basic helix-loop-helix leucine zipper transcription factor the MiT/TFE family (Rehli *et al*, 1999). TFEB binds a promoter motif responsible for coordinating the expression of lysosomal genes; identified as the coordinated lysosomal expression and regulation (CLEAR) element (Sardiello *et al*, 2009). In addition, TFEB regulates autophagy through enhancing autophagosome formation and autophagosome-lysosome fusion (Settembre *et al*, 2011). Stress conditions, such as starvation or exposure to ER stress-inducing chemicals, result in translocation of TFEB to the nucleus, where it promotes transcription of its target genes (Sardiello *et al*., 2009; Settembre *et al*., 2011). TFEB nuclear translocation is associated with its phosphorylation state (Settembre *et al*., 2011). The increase of Ca^2+^ levels leads to the activation of the phosphatase calcineurin, which dephosphorylates TFEB, resulting in TFEB nuclear translocation and the transcription of target genes (Medina *et al*, 2015). Enhancing the ALP through TFEB overexpression has marked beneficial effects in ameliorating amyloidogenesis in neurodegenerative diseases (Decressac *et al*., 2013; Sardiello *et al*., 2009; Tsunemi *et al*., 2012).

We demonstrate here that PERK activation can mitigate amyloidogenesis through promoting ALP. SB202190, previously identified as a p38 MAPK inhibitor, increases TFEB-ALP *via* Ca^2+^-dependent calcineurin activation independent of p38 MAPK inhibition (Yang *et al*, 2020). We describe a new function of SB202190, demonstrating that this molecule can activate PERK through mtROS production leading to the release of Ca^2+^ from the ER in human neuroblastoma cells. SB202190-induced PERK activation enhances TFEB nuclear translocation, leading to activation of the ALP. Finally, we demonstrate that PERK activation, as a novel target of SB202190, promotes the degradation of APP or α-syn accumulation *via* ALP in SH-SY5Y cells. These studies confirm the feasibility of PERK pathway targeting as a therapeutic approach for neurodegenerative diseases.

## Results

### SB202190 activates the PERK/eIF2α/ATF4 pathway

Our previous data have suggested that PERK activation can promote calcineurin-dependent TFEB nuclear translocation, and subsequent increase of autophagy and lysosomal-related gene expression (Kim *et al*, 2018). A recent study has found that only SB202190, among several known p38 MAPK inhibitors, can promote autophagy and lysosomal biogenesis dependent on calcineurin rather than *via* the inhibition of p38 MAPK (Yang *et al*., 2020). Therefore, to investigate the mechanisms by which SB202190 can induce autophagy and lysosomal biogenesis, and to determine the role of Ca^2+^ and PERK signaling underlying the response, we first assessed whether SB202190 activates PERK in various cells. SB202190 increased PERK activation in a dose-dependent manner in HEK293 cells (Fig 1A). We also confirmed that the activation of PERK by SB202190 was independent of p38 MAPK, using siRNA targeting p38 MAPK (si-p38 MAPK). PERK activation by SB202190 or thapsigargin (Tg) was similar in both the scrambled control RNA (scRNA)-transfected and si-p38 MAPK-transfected SH-SY5Y cells (Appendix Fig S1). In addition, PERK as well as its downstream targets, eIF2α and ATF4, were activated by SB202190 treatment, similar to the effect elicited by Tg, an inhibitor of the ER Ca^2+^-ATPase, used as a positive control (Fig 1B). To investigate the role of SB202190 in the activation of the PERK pathway, HEK293 cells were pretreated with a PERK inhibitor (GSK2606414). The inhibition of PERK with GSK2606414 attenuated the activation of eIF2α and ATF4 in response to SB202190 (Fig 1B). To investigate the effects of PERK activation in neuronal cells, the human neuroblastoma cell line SH-SY5Y was treated with SB202190. As expected, we found that SB202190 activated the PERK pathway in control siRNA-transfected cells, but not in SH-SY5Y cells subjected to siRNA-mediated PERK knockdown (Fig 1C). Consistent with observations in SH-SY5Y cells, treatment with SB202190 induced the activation of the PERK pathway in MEF cells isolated from *Perk*^*+/+*^ mice, but not *Perk*^*-/-*^ mice (Fig 1D). To assess whether SB202190 can activate the other UPR branches (i.e., IRE1α and ATF6), we analyzed the levels of Xbp1 splicing and GRP78 expression in SB202190-treated SH-SY5Y cells. These events were not affected by SB202190 treatment but were responsive to Tg treatment (Fig 1E). The activation of the PERK pathway by SB202190 was not affected by genetic deficiency of IRE1α or ATF6 (Fig 1F and 1G). These results demonstrate that SB202190 can selectively induce activation of the PERK branch of the UPR, but not IRE1α or ATF6.

**Figure 1.**
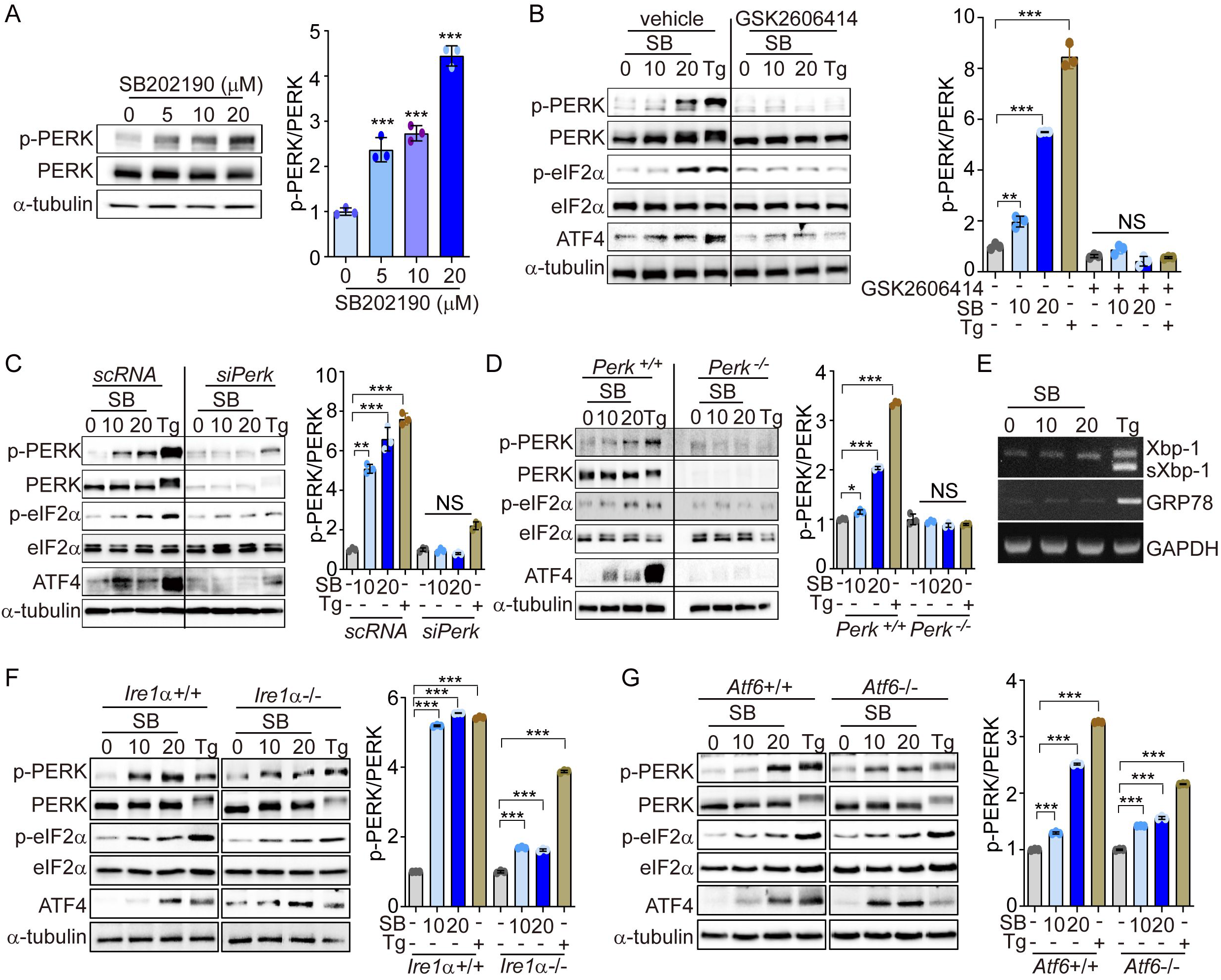
SB202190 activates the PERK/eIF2α/ATF4 pathway. (A) HEK293 cells were treated with SB202190 (0, 5, 10, and 20 μM) for 6 h. PERK phosphorylation was determined by western blotting. Quantification of p-PERK is shown in the right panel. (B) HEK293 cells were incubated with SB202190 (10 and 20 μM) for 6 h after pretreatment with or without the PERK inhibitor, GSK2606414 (1 μM) for 1 h. Thapsigargin (Tg, 2 μM) was used as a positive control. Cell lysates were used for western blotting analysis for p-PERK, PERK, p-eIF2α, eIF2α, and ATF4. (C) For knockdown of *Perk*, SH-SY5Y cells were transfected with control siRNA (scRNA) or si*Perk* for 48 h and then treated with different doses (10 and 20 μM) of SB202190 (SB) or Tg (2 μM) for 6 h. Cell lysates were measured for PERK activation by western blot using the indicated antibodies. (D) *Perk*^*+/+*^ and *Perk*^*-/-*^ MEFs were treated with SB202190 (10 and 20 μM) for 6 h or Tg (2 μM). The levels of p-PERK, PERK, p-eIF2α, eIF2α, and ATF4 were measured by western blotting. (E) SH-SY5Y cells were treated with SB202190 (10 and 20 μM) or Tg (2 μM) for 6 h. *Xbp-1* splicing and *Grp78* expression were detected by RT-PCR. (F, G) Hepatocytes isolated from *Ire1α*^*+/+*^, *Ire1α*^*-/-*^ (F), or *Atf6α*^*+/+*^, and *Atf6α* ^*-/-*^ (G) mice were treated with SB202190 (10 and 20 μM) for 6 h to assess the levels of PERK phosphorylation and ATF4 expression by western blotting. Quantification of p-PERK is shown in the right panel (A, B, C, D, F, and G). Data are mean ± SD (*n*=3); \**p*<0.05, \*\**p*<0.01, and \*\*\**p*<0.001.

### PERK activation with SB202190 is induced by mtROS production

Low levels of mtROS can selectively activate the PERK/eIF2α/ATF4 axis to preserve cellular homeostasis (Chartier-Harlin *et al*, 2004; Chen *et al*, 2019; Joe *et al*, 2018; Kim *et al*., 2018). Hughes *et al*., suggested that the modulation of the PERK pathway can protect against neurodegeneration (Hughes & Mallucci, 2019). Thus, we first investigated whether SB202190-induced PERK activation was regulated by mtROS in human neuronal cells. The levels of mtROS after SB202190 treatment were detected using the mitochondrial superoxide indicator MitoSOX in SH-SY5Y cells. In SH-SY5Y cells, treatment with SB202190 enhanced mtROS levels as analyzed by confocal microscopy (Fig 2A) and FACS (Fig 2B). Conversely, SB202190 reduced mtROS levels were diminished in cells treated with MitoTEMPO, an mtROS scavenger (Fig 2A and 2B). In accordance with observations in SH-SY5Y cells, we confirmed that SB202190 enhanced the generation of mtROS in HEK293 cells (Appendix Fig S2A). To determine whether SB202190 treatment activates the PERK pathway *via* mtROS, the levels of phosphorylation of PERK, eIF2α, and ATF4 were measured in MitoTEMPO-treated SH-SY5Y and HEK293 cells. Scavenging mtROS efficiently reduced PERK pathway activation in SB202190-treated cells (Fig 2C and Appendix Fig S2B). These data demonstrate that SB202190 treatment is effective in activating PERK *via* mtROS generation, specifically in neurons.

**Figure 2.**
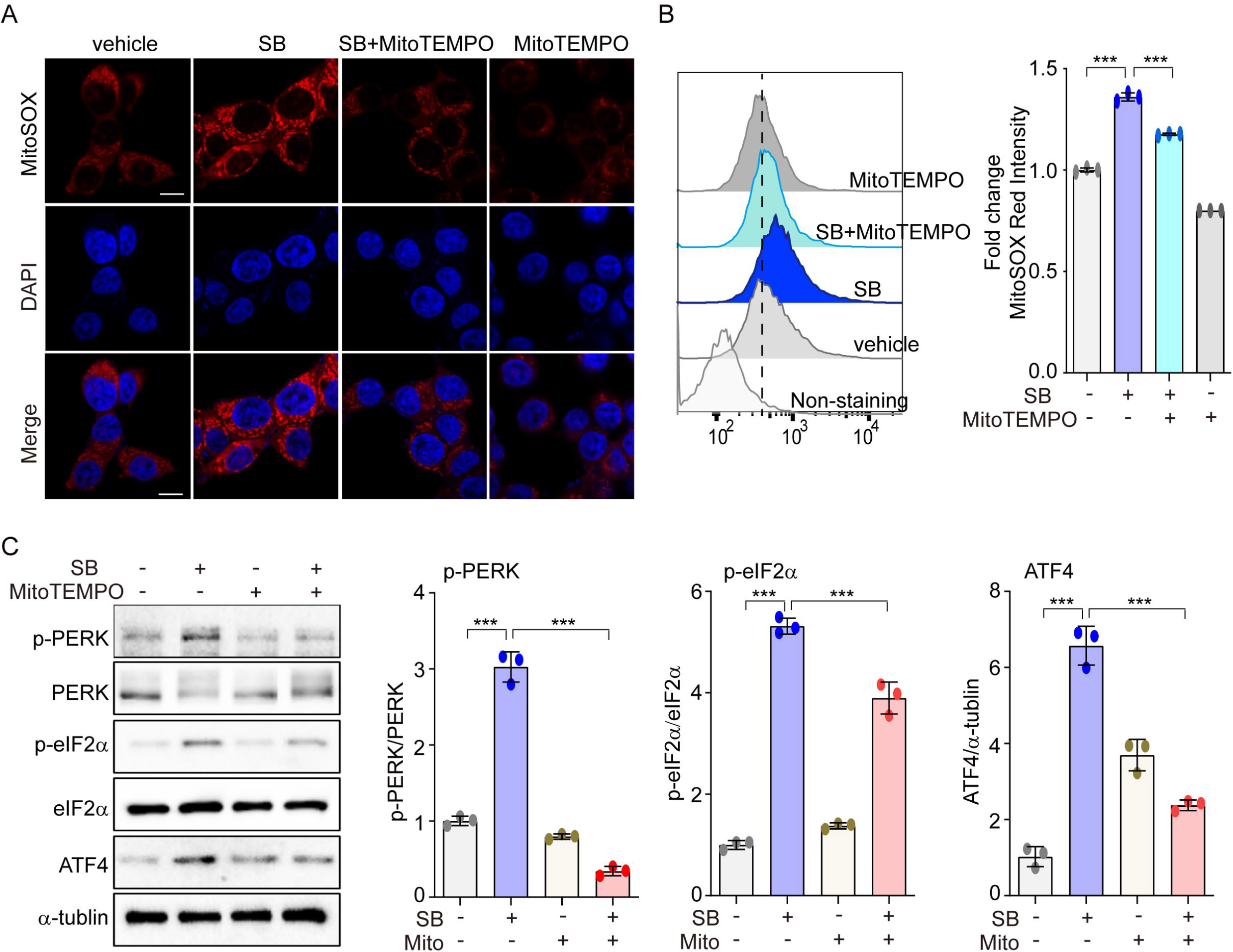
PERK activation with SB202190 is induced by ROS production. (A-C) SH-SY5Y cells were treated with 20 μM SB202190 for 3 h in the presence or absence of MitoTEMPO (100 nM, 1 h). (A) Cells were stained with MitoSOX (red), and then the nuclei were stained with DAPI (blue). Images were acquired by confocal microscopy. Scale bar, 10 μm. (B) mtROS was measured by flow cytometry. Fold changes in MitoSOX intensity are indicated at right. Data are mean ± SD (*n*=3); \*\*\**p*<0.001. (C) Western blotting was performed to detect the phosphorylation of PERK, eIF2α, and ATF4.

### PERK is required for TFEB nuclear translocation by SB202190 in SH-SY5Y cells

PERK is involved in translocation of TFEB into the nucleus to induce the autophagy-lysosomal pathway (Kim *et al*., 2018). In neuronal cells, the activation of ALP by increase of TFEB nuclear translocation is critical to ameliorate amyloidogenesis. We investigated whether PERK affects TFEB nuclear translocation in SB202190-treated SH-SY5Y cells. SB202190 induced a dose-dependent increase in TFEB nuclear translocation, as did starvation, compared to serum-fed cells (Fig 3A). In addition, cells expressing TFEB-GFP revealed that TFEB redistributes from the cytosol to the nucleus after SB202190 treatment (Fig 3B). The inhibition of PERK with GSK2606414 prevented the nuclear translocation of TFEB-GFP by SB202190, indicating that SB202190-regulated TFEB activity is mediated by PERK (Fig 3B). In agreement with observations in PERK inhibitor-treated cells, genetic deficiency of PERK abolished SB202190-induced TFEB nuclear translocation (Fig 3C). In contrast to the requirement for PERK in SB202190-induced TFEB activity; inhibition of mTOR with Torin1 treatment, promote TFEB nuclear translocation in a manner independent of PERK. Conversely, Tg treatment was found to inhibit TFEB nuclear translocation consistent with a previous report (Medina *et al*., 2015). These results demonstrate that TFEB nuclear translocation is required for PERK activation in human neuroblastoma cell lines in response to SB202190.

**Figure 3.**
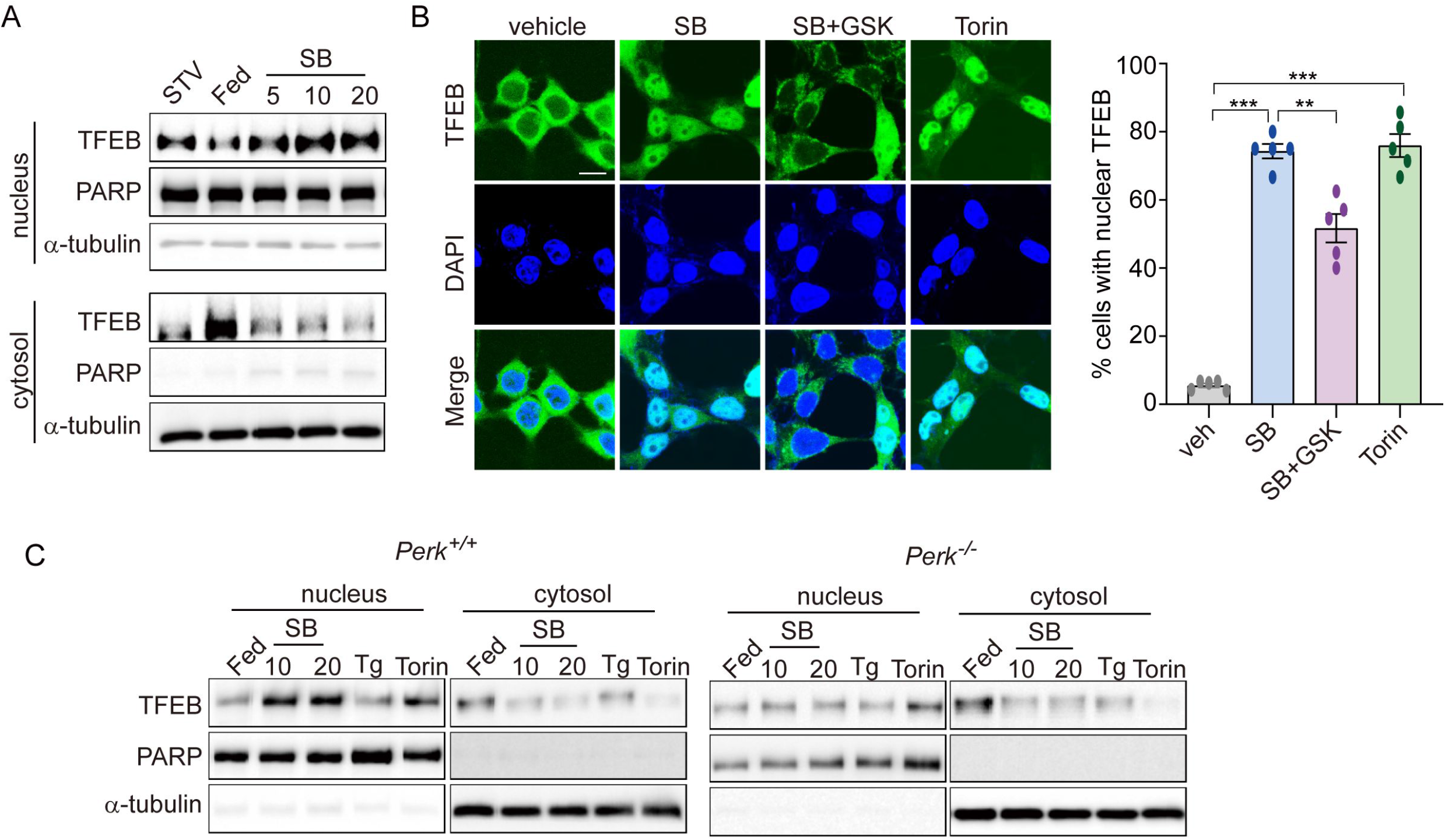
PERK is required for TFEB nuclear translocation by SB202190. (A) SH-SY5Y cells were treated with SB202190 at the indicated concentrations (5, 10, and 20 μM) for 3 h and subjected to nuclear and cytosolic fractionation. Resulting fractions were then detected with antibody against TFEB. Starvation (STV) or HBSS was used as positive control or negative control, respectively. PARP and α-tubulin were used as nuclear and cytosolic markers, respectively. (B) TFEB-EGFP-transfected SH-SY5Y cells were treated with 20 μM SB202190 in the presence or absence of the PERK inhibitor (GSK2606414, GSK). Torin-1, an mTOR inhibitor, was used as a positive control. The fluorescence of TFEB was visualized by confocal microscopy (*left*). Scale bar: 10 μm. Cells were evaluated to calculate the percentage of cells showing nuclear TFEB localization. *n* > 20 cells per condition. Data are mean ± SD; \*\**p*<0.01 and \*\*\**p*<0.001. (C) *Perk*^*+/+*^ and *Perk*^*-/-*^ MEFs were treated with SB202190 (10 and 20 μM) for 6 h or Tg (2 μM). Torin-1 (2 μM) treatment was used as a positive control. Cells were detected with TFEB antibody in the nuclear and cytosol fraction by western blotting.

### The PERK-Ca^2+^-calcineurin pathway is required for SB202190-induced TFEB nuclear translocation

We next investigated whether SB202190-induced PERK activation triggers Ca^2+^ release from the ER. In our previous study, we demonstrated that PERK activation induces the increase of cytosolic Ca^2+^ (Kim *et al*., 2018). Consistent with that finding, SB202190-induced PERK activation increased cytosolic Ca^2+^ in a dose-dependent manner (Fig 4A). Conversely, the PERK inhibitor GSK2606414 inhibited the cytosolic Ca^2+^ increase caused by SB202190 (Fig 4B). The increase of cytosolic Ca^2+^ in SB202190-treated cells was verified using Fluo-4AM (Fig 4C). In cells loaded with Fluo-4AM in the presence of SB202190, Ca^2+^ levels were decreased by treatment with GSK2606414 (Fig 4C). These results suggest that the SB202190-induced increase in cytosolic Ca^2+^ is mediated *via* PERK activation. TFEB activity is regulated by phosphorylation (Settembre *et al*, 2012), which retains inactive TFEB in the cytoplasm. In contrast, when dephosphorylated by the phosphatase calcineurin, TFEB translocates to the nucleus to activate transcriptional target genes (Medina *et al*., 2015). We have also reported that PERK activation induced Ca^2+^-calcineurin-dependent TFEB regulation (Kim *et al*., 2018). In this study, to explore the mechanisms of SB202190-induced TFEB nuclear translocation, SH-SY5Y cells, in the presence of SB202190, were co-incubated with the calcineurin inhibitors, FK506, or cyclosporin A (CsA). SB202190-induced TFEB nuclear translocation was suppressed by FK506 (Fig 4D) and CsA (Fig 4E), suggesting that calcineurin is involved in SB202190-induced TFEB nuclear translocation. We confirmed that calcineurin inhibitors block SB202190-induced TFEB nuclear translocation in SH-SY5Y cells expressing TFEB-GFP (Fig 4E). These results, taken together, implicate an important role of the PERK-Ca^2+^-calcineurin pathway in SB202190-induced TFEB activation in human neuroblastoma cells.

**Figure 4.**
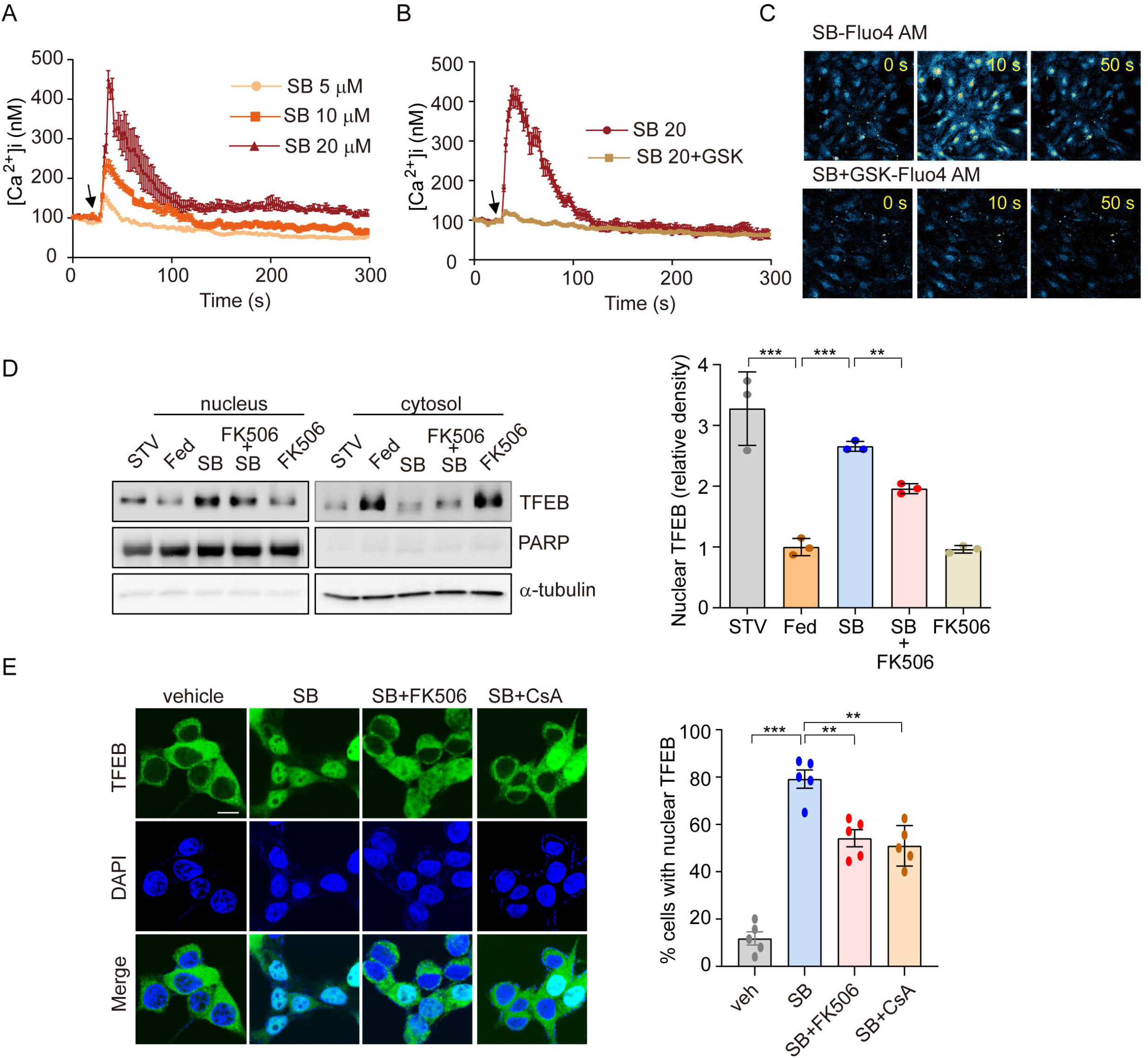
PERK-Ca^2+^-calcineurin pathway is required for SB202190-induced TFEB nuclear translocation. (A-C) The change of [Ca^2+^] in MEF cells was measured using confocal microscopy after loading with Fluo-4 AM. (A) Arrow indicates the time point at which SB202190 (SB) was added. The data represent mean ± SD from 3 independent experiments. (B, C) MEF cells were preincubated for 30 min with the PERK inhibitor GSK2606414 (1 μM). Arrow indicates the time point at which 20 μM SB202190 was added. The data represent mean ± SD from 3 independent experiments. (D) SH-SY5Y cells were pretreated with FK506 (10 μM) for 30 min and then treated with SB202190 (20 μM) for another 3 h. Measurement of TFEB activation was performed by western blotting of nuclear and cytoplasmic extracts. TFEB expression in the nucleus and cytoplasm was normalized to PARP and α-tubulin, respectively. (E) TFEB-GFP-transfected SH-SY5Y cells were treated with SB202190 (20 μM) for 6 h in the presence or absence of calcineurin inhibitors, FK506 (10 μM) and Cyclosporin A (CsA, 20μM). Representative images were detected by confocal microscopy (*left*). Quantification of nuclear translocation of TFEB-GFP (right). *n* > 20 cells per condition. Data represent mean ± SD; \*\**p*<0.01 and \*\*\**p*<0.001.

### PERK activation by SB202190 facilitates autophagy and lysosome biogenesis *via* TFEB activation

Coordination of the multiple steps in the ALP requires the existence of TFEB as a master regulator (Martini-Stoica *et al*., 2016). The main function of TFEB was identified in coordinating autophagy through regulating autophagosome formation and autophagosome-lysosome fusion (Settembre *et al*., 2011). TFEB also promotes cellular clearance through lysosomal exocytosis, a process mediated by activation of the lysosomal Ca^2+^ channel MCOLN1 (Medina *et al*, 2011). To investigate whether SB202190-induced TFEB activation enhances the activation of autophagy-lysosomal axis, we first measured LC3B-II and p62 levels. Consistent with a previous study (Yang *et al*., 2020), SB202190 increases the levels of LC3B-II and p62 in a dose-dependent manner (Fig 5A). To further investigate autophagy flux, we analyzed autophagosome (yellow puncta) and autolysosome (red puncta) in SH-SY5Y cells transfected with the mCherry-EGFP-LC3 reporter. Autolysosome levels were increased by SB202190, and were further enhanced following chloroquine (CQ) treatment (Fig 5B). Moreover, LC3B-II conversion in the presence of CQ was greater than that observed in SB202190-treated cells without chloroquine, suggesting that SB202190 stimulates autophagic flux (Fig 5C). Next, we examined whether the effect of SB202190 on lysosomal biogenesis is dependent on PERK activation. First, we observed significant and dose-dependent increases in mRNA levels of lysosomal genes, including cathepsin D, cathepsin B, LAMP1, MCOLN1, and TPP1, in SB202190-treated SH-SY5Y cells (Appendix Fig S3A). Consistent with the PERK-dependency of TFEB activation in response to SB202190 treatment (Fig 3), we observed that the expression levels of lysosomal genes were increased by SB202190 treatment in *Perk*^*+/+*^ cells, but not in *Perk*^*-/-*^ cells (Fig 5D). In contrast, the ability of SB202190 to increase the mRNA expression of lysosomal genes was not affected by genetic deficiency of IRE1α and ATF6α (Appendix Fig S3B and S3C). In addition, deficiency of TFEB clearly suppressed SB202190-increased levels of LC3B-II and p62 (Fig 5E), and lysosomal biogenesis, using LysoTracker staining and real time PCR (Fig 5F-5H). Overall, these results revealed that PERK activation by SB202190 requires the ALP through TFEB activation

**Figure 5.**
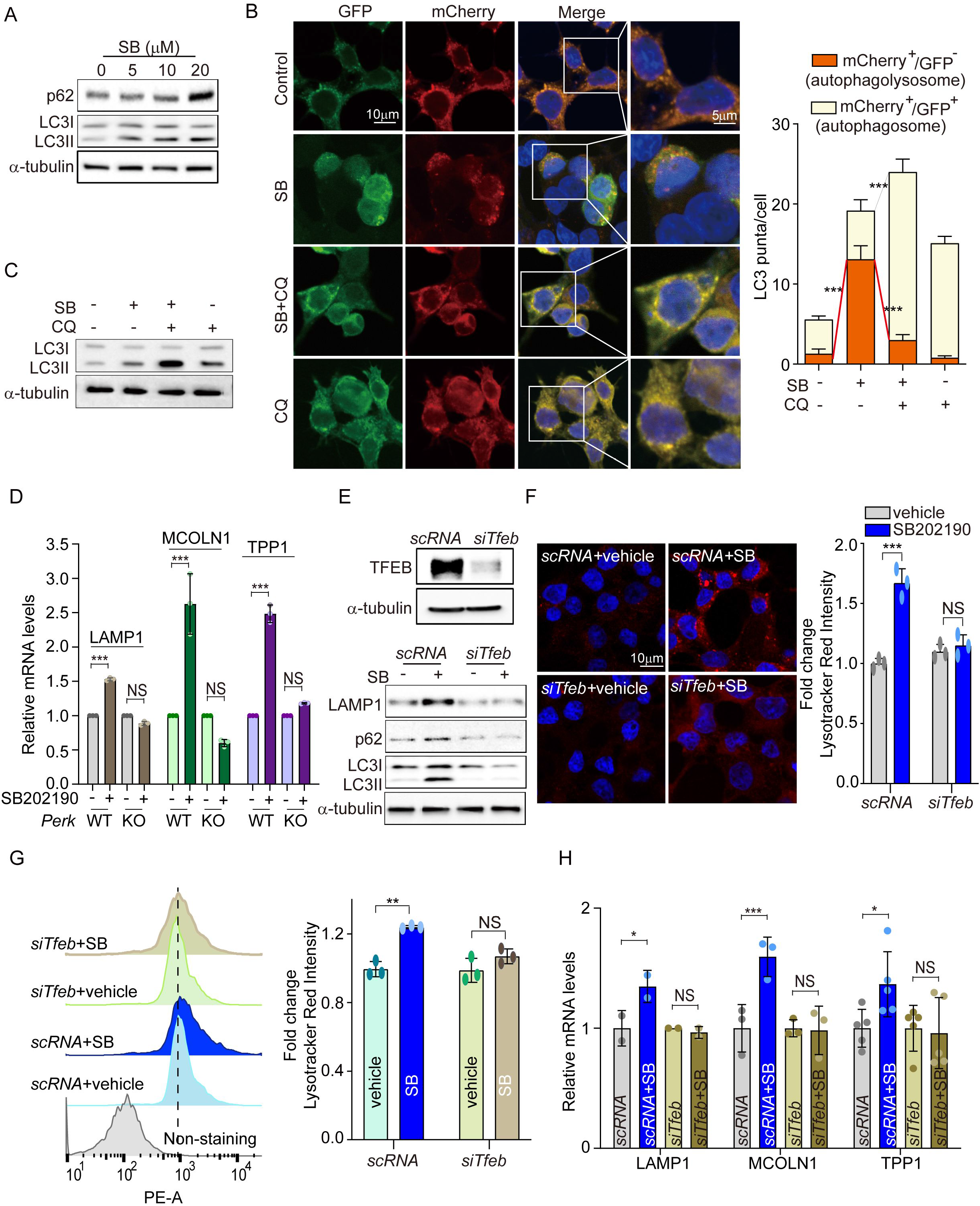
PERK activation by SB202190 facilitates autophagy and lysosome biogenesis via TFEB activation. (A) HEK293 cells were treated for 6 h with SB202190 (5, 10, and 20 μM) and subjected to western blotting by antibodies against p62 and LC3B. (B) SH-SY5Y cells were transiently transfected with mCherry-GFP-LC3 for 48 h and subsequently pretreated with chloroquine (CQ, 10 μM) for 1 h, and then treated with SB (20 μM) for 6 h. Cells were observed for fluorescence of both GFP and mCherry using confocal microscopy. The number of autolysosomes (GFP-RFP^+^) and autophagosomes (GFP^+^RFP^+^) per cell in each condition were quantified. Data represent mean ± SD; \*\*\**p*<0.001. (C) SH-SY5Y cells were treated with CQ (10 μM) for 1 h before SB202190 (20 μM) for 6 h. The levels of LC3B-II conversion were analyzed by western blotting. (D) To check of PERK dependence in lysosome biogenesis, *Perk*^*+/+*^ and *Perk*^*-/-*^ MEFs were treated with SB202190 (20 μM) for 6 h. Lysosomal genes, LAMP1, MCOLN1, and TPP1, were measured by qRT-PCR. Data represent mean ± SD; \*\*\**p*<0.001, NS, not significant. (E-H) SH-SY5Y cells were transfected with *siTfeb* for 48 h and then treated with SB202190 (20 μM) for 6 h. (E) Cells were subjected to western blotting by using antibodies against TFEB (*upper*), LAMP1, p62, and LC3B (*lower*). (F) Samples were stained with Lysotracker Red. Representative image was obtained by confocal microscopy (*left*). Scale bar, 10 μm. Quantification of lysosome intensity was determined by counting red puncta (r*ight*). Data represent mean ± SD; \*\*\**p*<0.001, NS, not significant. (G) LysoTracker fluorescence was measured by flow cytometry. Fold changes in LysoTracker intensity are indicated at the right and are presented as mean ± SD (n=3); \*\**p*<0.01, NS, not significant. (H) lysosomal genes, LAMP1, MCOLN1, and TPP1, were measured by qRT-PCR. Data are represented as mean ± SD; \**p*<0.05, \*\*\**p*<0.001, NS, not significant.

### PERK activation by SB202190 reduces the aggregation of APP and α-syn accumulation through TFEB-ALP activation in SH-SY5Y cells

PERK activation enhances the ALP *via* TFEB activation to degrade misfolded proteins that accumulate in neurodegenerative disorders (Ganz *et al*, 2020; Song *et al*, 2020). A PERK activator was shown to act as a potent agent that reduces toxicity and extends survival in murine models of HD (Ganz *et al*., 2020). Accumulations of APP and α-syn have been observed in AD and PD, respectively (Martini-Stoica *et al*., 2016). To explore whether SB202190-induced PERK activation attenuates amyloidogenesis, SH-SY5Y cells were transfected with plasmids encoding APP 695 Swedish and Indiana mutation (APP^Swe/Ind^) or α-synuclein A53T mutation (α-Syn-A53T). The aggregated proteins of full length-APP (FL-APP) or α-syn were increased by SB202190 treatment and conversely were inhibited by the PERK inhibitor, GSK2606414 (Fig 6A and Appendix Fig S4A). Consistent with the function of SB202190 in degradation of neurotoxic proteins, LC3B-II conversion was increased after SB202190 treatment but not in the presence of PERK inhibitor (Fig 6B and Appendix Fig S4B), suggesting that autophagy processing may mediate the SB202190-induced decrease of APP and α-syn. In this study, we demonstrated that PERK activation by SB202910 treatment in SH-SY5Y cells was mediated by mtROS production (Fig 2). Thus, to confirm the role of mtROS in the degradation of aggregated mutant amyloid protein, APP^Swe/Ind^ transfected SH-SY5Y cells were treated with MitoTEMPO. As expected, the decrease of mutant APP accumulation was reversed by MitoTEMPO treatment. As shown in Fig 5, ALP activation by SB202190 in SH-SY5Y cells was caused by TFEB activity. Therefore, to further investigate whether the effect of SB202190 on degradation of accumulated mutant APP was dependent on TFEB, we performed co-transfection of SH-SY5Y cells with si*Tfeb* and APP^Swe/Ind^. TFEB deficient cells failed to degrade FL-APP after SB202190 treatment, indicating that TFEB activation by SB202190 is essential to activate ALP for the reduction of neurotoxic proteins in SH-SY5Y cells (Fig 6C and 6D). Taken together, our results demonstrate that SB202190-induced ALP activation for degradation of mutant APP and α-syn accumulation requires PERK-dependent TFEB activation.

**Figure 6.**
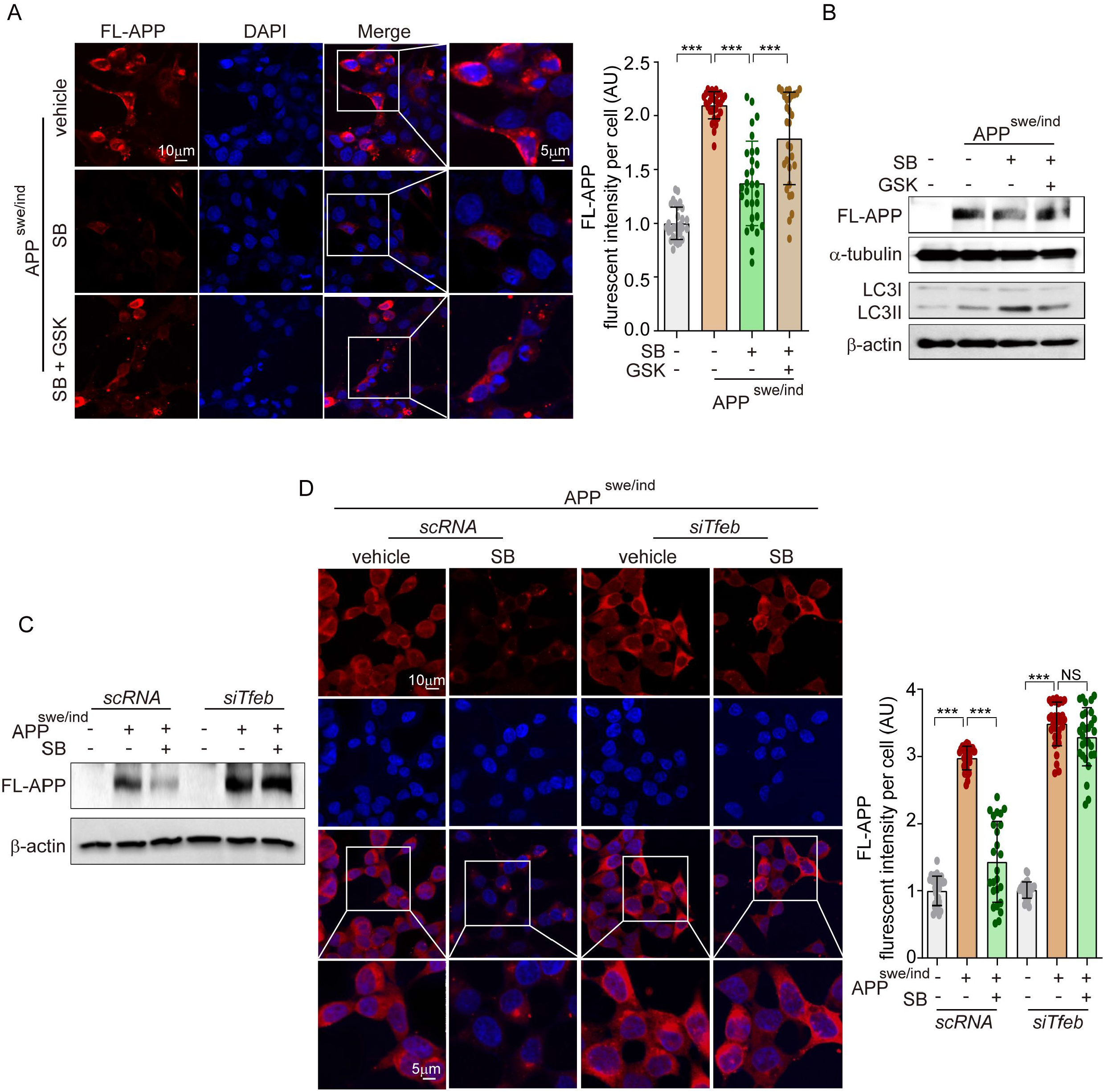
PERK activation by SB202190 reduces the aggregation of APP accumulation through TFEB-ALP activation in SH-SY5Y cells. (A-B) SH-SY5Y cells were transfected with pCAX-APP-Swe/Ind (APP^swe/ind^) for 48 h. Under this condition, cells were treated with the PERK inhibitor, GSK2606414 (1 μM, 1 h), before SB202190 (20 μM, 12 h) treatment. (A) Cells were stained with DAPI and immunostained with anti-FL-APP antibody. Representative image of FL-APP was observed by confocal microscopy (*left*) and quantification of FL-APP intensity (*right*). Data represent mean ± SD; \*\*\**p*<0.001. (B) Western blotting was performed to detect the expression levels of FL-APP and LC3B. (C, D) SH-SY5Y cells were co-transfected with pCAX-APP-Swe/Ind (APP^swe/ind^) and si*Tfeb* for 48 h and were treated with SB202190 (20 μM, 12 h). (C) Cells were detected to antibody against FL-APP by western blotting. (D) Cells were stained with antibody against FL-APP and DAPI. Representative image of FL-APP was obtained by confocal microscopy (*left*) and quantification of FL-APP intensity (*right*). Data represent as mean ± SD; \*\*\**p*<0.001 and not significant (NS).

## Discussion

This study identifies a novel effect of SB202190 in promoting PERK activation, independent of p38 MAPK inhibition, in the cellular response to mitigate amyloidogenesis in human neuroblastoma cells. We demonstrate that SB202190-induced PERK activation activates TFEB nuclear translocation, leading to induction of the ALP. The ALP is involved in the degradation of misfolded proteins that accumulate in many neurodegenerative diseases such as AD, PD, and HD (Martini-Stoica *et al*., 2016). Thus, enhancing the activity of ALP in neurodegenerative diseases has been extensively studied as a strategy for therapeutic intervention (Menzies *et al*., 2015). Interestingly, only SB202190, among several other p38 MAPK inhibitors, can enhance TFEB-mediated ALP activation (Yang *et al*., 2020). SB202190 triggers the release of Ca^2+^ from the ER and subsequent calcineurin activation, leading to TFEB-induced ALP activation, apart from its known function as a p38 MAPK inhibitor (Yang *et al*., 2020). Our data also show that SB202190 promotes TFEB nuclear translocation in SH-SY5Y cells. We further found that SB202190 specifically induces the activation of PERK among the three known ER sensors, PERK, IRE1α and ATF6. In animal models of neurodegenerative disease and in human post-mortem tissue from many neurodegenerative diseases (Hughes & Mallucci, 2019; Moreno *et al*, 2012), the PERK pathway has been associated with disease progression. Genetic and pharmacological modulation of the PERK pathway is the focus of new therapeutic strategies of neurodegenerative diseases (Hughes & Mallucci, 2019; Imran & Mahmood, 2011; Radford *et al*, 2015). Therefore, our findings for SB202190-induced PERK activation in neuroblastoma cells may provide insight into strategies to resolve pathological amyloidogenesis.

Our results also indicate that PERK activation by SB202190 is mediated by mtROS, resulting in the increase of cytosolic Ca^2+^ and the activation of calcineurin. In our previous studies, we found that a mild increase of mtROS levels leads to PERK activation, Ca^2+^ release from the ER, and calcineurin-dependent TFEB activation in hepatocytes (Kim *et al*, 2017; Kim *et al*., 2018). Moreover, calcineurin plays a detrimental role in damaged neurons. Notably, the cytoprotective effects of calcineurin in acute brain injuries are PERK-dependent (Chen *et al*., 2016). Thus, our study suggests that the protective role of PERK-calcineurin in neurons is due to an increase TFEB nuclear translocation. Calcineurin dephosphorylates TFEB, leading to increased autophagy induction and lysosomal biogenesis (Medina *et al*., 2015). We demonstrate that SB202190 increases autophagy-related genes and autophagolysosome formation. In addition, our data show that SB202190-induced ALP activation is dependent on PERK but not IRE1α and ATF6 in human neuroblastoma cells. To verify the amelioration of amyloidogenesis *via* SB202190-induced ALP activation, SH-SY5Y cells were transfected with mutant FL-APP or α-syn as *in vitro* models of AD or PD, respectively (Chartier-Harlin *et al*., 2004; Song *et al*., 2020; Zhang *et al*, 2011). Accumulation of FL-APP or α-syn in SH-SY5Y cells was mitigated with SB202190 treatment. In particular, the decrease of amyloidogenesis with SB202190 required PERK activation. Thus, we also demonstrated that SB202190-induced PERK activation ameliorates amyloidogenesis in AD or PD models *via* mtROS production using the mitochondria-targeting antioxidant MitoTEMPO.

In summary, our data reveal a novel cytoprotective role for SB202190 as a PERK activator in human neuroblastoma cells. Our findings also suggest that SB202190 activates PERK *via* an increase of mtROS, leading to calcineurin activation. Subsequently, PERK activation by SB202190 leads to an increase in TFEB nuclear translocation through calcineurin-dependent dephosphorylation of TFEB. Finally, SB202190-PERK-TFEB activation attenuates amyloidogenesis *via* the increase of the ALP (see schema in Fig. 7). We anticipate that SB202190 and related compounds targeting PERK can be developed as therapeutics to ameliorate amyloidogenesis in neurodegenerative disorders.

**Figure 7.**
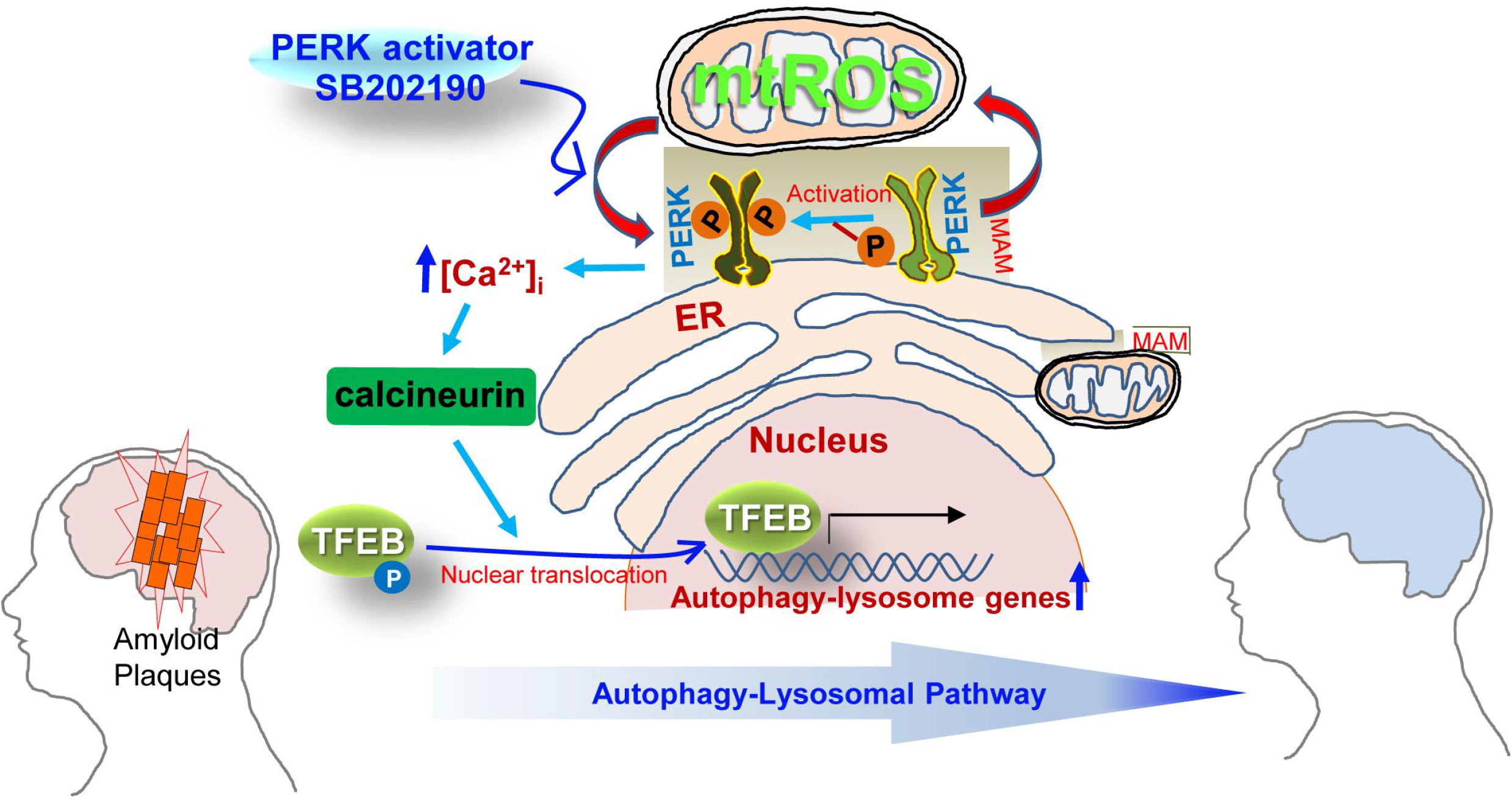
Schematic overview of the mechanisms by which SB202190 ameliorates amyloidogenesis *via* PERK activation. PERK, a tethering molecule of the mitochondrial-associated ER membrane (MAM), is activated by mitochondrial ROS (mtROS) in response to treatment with the PERK activator (SB202190). Activated PERK leads to increase in cytosolic Ca^2+^ levels and subsequently promotes the translocation of TEFB into the nucleus *via* the calcineurin-dependent dephosphorylation TFEB, which culminates in the increased transcription of autophagy-lysosome related genes. The increase of autophagy-lysosomal pathway (ALP) by PERK activation enhances the degradation of misfolded proteins that accumulate in neurodegenerative disorders. Therefore, the PERK-TFEB-ALP pathway activated by SB202190 suggests the novel target for ameliorating amyloidogenesis.

## Materials and Methods

### Reagents

SB202190, GSK2606414, thapsigargin (Tg), FK506, chloroquine (CQ), and cyclosporin A (CsA) were purchased from Sigma-Aldrich (St Louis, MO, USA). Torin 1 was from (Tocris-Biotechne, Minneapolis, MN.

### Cell lines

The human neuroblastoma cell line SH-SY5Y, and the human embryonic kidney cell line HEK293 were grown in Dulbecco’s Modified Eagle medium, DMEM (Gibco, Grand Island, USA), supplemented with 10% fetal bovine serum (Gibco, Melbourne, Australia) and 1% penicillin-streptomycin solution (Gibco, Grand Island, USA). *Perk*^*+/+*^ and *Perk*^*-/-*^ mouse embryonic fibroblasts (MEFs) were cultured in DMEM supplemented with 1% MEM non-essential amino acid (Gibco). *Ire1α*^*+/+*^, *Ire1α*^*-/-*^, *Atf6α*^*+/+*^, and *Atf6α*^*-/-*^ mouse hepatocytes, which were kindly provided by Dr. S. H. Back (University of Ulsan, Ulsan, Korea), were maintained in 199 medium, with 1% MEM non-essential amino acid solution. Cells were grown in humidified incubators, at 37°C with 5% CO_2_.

### Western blot

Total protein extracts from harvested-cells were prepared in RIPA buffer (Thermo Scientific, Waltham, MA, USA) containing phosphatase and protease inhibitors (Sigma-Aldrich), and the protein concentration was determined using the BCA protein assay kit (Pierce Biotechnology, Rockford, IL, USA) using bovine serum albumin (BSA) as standard. Aliquots of protein were boiled at 95°C in 2X Lamelli buffer (Bio-Rad, Hercules, CA, USA) for 5 min. Proteins were separated by SDS-PAGE and transferred to polyvinylidene difluoride membrane (Millipore, Burlington, MA, USA). The membrane was blocked with 5% nonfat milk (BD bioscience, San Jose, CA, USA) in phosphate buffered saline-Tween 20 (PBS-T), and then the membrane was immunoblotted with primary antibodies as follows: p-PERK (1:1000, Signalway Antibody, Baltimore, MD, USA), PERK (1:1000, Cell Signaling, Danvers, MA, USA), p-eIF2α (1:1000, Cell Signaling), eIF2α (1:1000, Cell Signaling), ATF4 (1:1000, Cell Signaling), TFEB (1:1000, Bethyl Laboratories, Montgomery, TX, USA), PARP (1:2000, Cell Signaling), LC3B (1:2000, Novus Biologicals, Centennial, CO, USA), p62 (1:10000), LAMP1 (1:1000, Abcam, Cambridge, MA, USA), FL-APP (1:2000, Sigma-Aldrich), α-synuclein (1:2000, Cell Signaling), α-tubulin (1:2000, Cell Signaling), and β-actin (1:2500, Thermo Scientific). This was followed by incubation in HRP-conjugated secondary antibodies. Immunoblots were detected using an ECL substrate (Pierce Biotechnology) and analyzed using an Azure Biosystems C300 analyzer (Azure Biosystems, Dublin, CA, USA).

### Measurement of mitochondrial ROS

SH-SY5Y and HEK293 cells were treated with SB202190 (20 μM) for 3 h in the presence or absence of 100 nM Mito-TEMPO (Sigma-Aldrich) for 1 h. After treatment, cells were stained with 5 μM MitoSOX Red (Invitrogen, Carlsbad, CA, USA) at 37°C for 30 min. For flow cytometry, cells were trypsinized and then washed three times with 1X PBS. mtROS were detected by using a FACSCanto flow cytometry system (BD Bioscience, CA, USA). Fluorescence data were analyzed with FlowJo software (C). For confocal microscope, cells were fixed with formalin solution for 30 min and then washed three times with 1X PBS. The cells were stained DAPI (Invitrogen). Samples were visualized by using an Olympus FV1200 confocal microscope (Olympus, Tokyo, Japan).

### RT-PCR and qRT-PCR

Total RNA was isolated from cells using by QIAzol Lysis reagent (QIAGEN, CA, USA). Reverse transcription of 2 μg RNA was performed using M-MLV reverse transcriptase (Promega, WI, USA). The PCR-based amplification was done using synthesized-cDNA. The following primers were human GAPDH (f-ccacccatggcaaattccatggca, r-tctagacggcaggtcaggtccacc), human Xbp1 (f-ccttgtagttgagaaccagg, r-ggggcttggtatatatgtgg), and human GRP78 (f-cgaggaggaggacaagaagg, r-ttgtttgcccacctccaata). To analyze real-time quantitative PCR (RT-qPCR), the synthesized cDNA was amplified with SYBR Green qPCR Master Mix on an ABI 7500 Fast Real-Time PCR System (Applied Biosystems, CA, USA), The following qRT-PCR primers were human GAPDH (f-caatgaccccttcatcctc, r-agcatcgccccacttgatt), human CTSB (f-agtggagaatggcacacccta, r-aagagccattgtcacccca), human CTSD (f-cttcgacaacctgatgcagc, r-tacttggagtctgtgccacc), human LAMP1 (f-cgtacctttccaacagcagc, r-cgctcacgttgtacttgtcc), human MCOLN1 (f-gagtgggtgcgacaagtttc, r-tgttctcttcccggaatgtc), and TPP1 (f-gatcccagctctcctcaatac, r-gccatttttgcaccgtgtg).

### Transfection

pEGFP-N1-TFEB, pBABE-puro mCherry-EGFP-LC3B, α-Syn-A53T, and pCAX-APP-Swe/Ind were purchased from Addgene (Watertown, MA, USA) and were transfected into cells *via* Lipofectamine™ 2000 (Invitrogen) in accordance with the manufacturer’s protocol. To knockdown the genes of *p38, Perk* and *Tfeb*, SH-SY5Y cells were transfected with scramble siRNA (scRNA) (Ambion, Austin, TX, USA), as the control siRNA, *p38* (Cell Signaling), *Perk* and *Tfeb* siRNA (Santa Cruz Biotechnology, CA, USA) using Lipofectamine™ 2000 (Invitrogen) method according to the manufacturer’s protocol. After 36 h, cells were treated with indicated drugs.

### Fluorescence cell imaging

For imaging of TFEB-GFP and mCherry-GFP-LC3, SH-SY5Y cells grown on coverslips were transfected with plasmids for 36 h and then were treated with indicated drugs. After treatment, cells were fixed with 10% formalin solution (Sigma-Aldrich) for 20 min at RT, followed by three times washes and DAPI staining. After additional three washes, the slides were mounted with mounting medium (Sigma-Aldrich). Cells were visualized using the Olympus FV1200 confocal microscope (Olympus).

### Immunofluorescence

To detect the FL-APP and α-synuclein using confocal microscope, SH-SY5Ys were incubated on coverslips and transfected with pCAX-APP695-Swe/Ind and α-syn-A53T, followed by the administration of SB202190, Mito-TEMPO, or PERK inhibitor. After treatment, the samples were fixed with 10% neutral-buffered formalin solution (Sigma Aldrich) for 20 min. The cells were subsequently permeabilized with 0.1% Triton X-100 and then blocked with 3% BSA for 30 min at RT. The cells were stained with anti-FL-APP (1:500, Sigma-Aldrich) or anti-α-synuclein (1:500, Cell Signaling) antibodies for overnight at 4 °C. Alexa-Fluor 594 anti-rabbit (1:500, Invitrogen) secondary antibody was added for 2 h, at RT. Then, the cells were washed with 1X PBS-T followed by DAPI staining. Images of the cells were obtained using an Olympus FV1200 confocal microscope (Olympus). The intensity of FL-APP and α-syn was analyzed by using ImageJ software.

### Subcellular fractionation

Nuclear and cytosolic fractions were prepared by using nuclear/cytosol fractionation kit (Biovision, CA, USA). Cells were harvested, and subcellular fractionation was performed according to the manufacturer’s instructions. Briefly, cell pellets were resuspended with cytosolic extraction buffer A (CEB-A) and CEB-B. After 10 min, the lysates were centrifuged at 4°C for 5 min at 16,000 X *g* in a microcentrifuge to obtain cytosolic protein containing supernatant. The remaining pellets were resuspended in nuclear extraction buffer (NEB) and vortexed for 15 s. This step was repeated every 10 min, for five times. Samples were centrifuged at 4°C for 10 min at 16,000 X *g* to acquire nuclear extraction. The purity of cytoplasmic and nuclear fractions was subjected to western blotting by anti-β-actin and PARP antibodies.

### Measurement of intracellular Ca^2+^ concentration

MEF cells were plated and cultured on confocal dishes. The cells were loaded with 5 μM fluo-4 AM (Invitrogen) in Hanks’ balanced salt solution (HBSS) with 1% bovine serum albumin at 37°C for 40 min as described previously (Kim *et al*., 2018). The cells were washed three times with HBSS. Changes of intracellular Ca^2+^ concentration ([Ca^2+^]_i_) were determined at 488 nm excitation/530 nm emission using an air-cooled argon laser system. The emitted fluorescence at 530 nm was collected using a photomultiplier. The image was scanned using a confocal microscope (Nikon, Japan). For the calculation of [Ca^2+^]_i_, the method of Tsien *et al*. (Tsien *et al*, 1982) was used with the following equation: [Ca^2+^]_i_ = Kd(F − Fmin)/(Fmax − F), where Kd is 345 nM for fluo-4 and F is the observed fluorescence level. Each tracing was calibrated for the maximal intensity (Fmax) by addition of ionomycin (8 μM) and for the minimal intensity (Fmin) by addition of EGTA (50 mM) at the end of each measurement.

### Lysosome quantification

The lysosomal contents were assessed using LysoTracker Red (Invitrogen) following manufacturer’s instructions. After treatment, cells in fresh medium were stained with LysoTracker Red for 1 h followed by extensive washing and then transferred into tubes for quantification of lysosome fluorescence using a FACSCanto flow cytometry system (BD Bioscience). Data were analyzed by FlowJo software (Tree Star). For measurement of lysosome intensity using confocal image, the stained cells were fixed with 10% formalin solution followed by DAPI staining. Samples expressing fluorescence were identified by Olympus FV1200 confocal microscope.

### Statistical Analysis

Data were analyzed with Prism (GraphPad Software, San Diego, CA, USA). All values are presented as mean ± SD. Statistical analyses were performed using Student’s *t*-test or one-way ANOVA with Tukey *post hoc* tests.

## Data availability

All relevant data are presented within the paper. Expanded View for this article is available online.

## Acknowledgements

This work was supported by the Priority Research Centers Program through the National Research Foundation of Korea (NRF) funded by the Ministry of Education (2014R1A6A1030318, NRF-2020R1A2C1009192) to H.T.C., NRF-2020R1A2C1006470 to Y. J

## Author contributions

Conceptualization: HTC, YJ; Data curation: JP, YJ; Formal analysis: MD, JP, YC, S-YR, T-HTN, JHG; Investigation: B-SK, JWP, U-HK, SWR, Y-JS; Project administration: YJ, HTC; Resources: JWP, U-HK, YJ, HTC; Writing, review and editing: B-SK, JWP, SWR, Y-JS, YJ, HTC.

## Figure Legends

**Appendix Figure S1. PERK activation with SB202190 is independent of p38 MAPK-related to Figure 1**.

(A, B) SH-SY5Y cells were transfected with control siRNA (scRNA) or si-p38 MAPK for 48 h and then treated with 20 μM SB202190 for 6 h or 1 μM Tg for 30 min. Cell lysates were evaluated for p38 MAPK knockdown by western blot (A) and detected with antibodies against p-PERK, PERK, p-eIF2α, eIF2α, and ATF4 by western blotting (B).

**Appendix Figure S2. PERK activation with SB202190 is induced by ROS production-related to Figure 2**.

(A, B) HEK293 cells were pretreated with MitoTEMPO (100 nM) for 1 h and then treated with 20 μM SB202190 for 3 h. (A) MitoSOX fluorescence was analyzed by flow cytometry (*left*), and quantification of fluorescence was normalized (*right*). (B) The activation of PERK was analyzed by western blotting using the indicated antibodies.

**Appendix Figure S3. PERK activation by SB202190 facilitates autophagy and lysosome biogenesis *via* TFEB activation related to Figure 5**.

(A) SH-SY5Y cells were treated with SB202190 (5, 10, and 20 μM) for 6 h, and then lysosomal genes were analyzed by qRT-PCR. Data represent mean ± SD; \*\*\**p*<0.001. (B, C) Hepatocytes from *Ire1α*^*+/+*^, *Ire1α*^*-/-*^ (B), *Atf6α*^*+/+*^, and *Atf6α*^*-/-*^ (C) mice were treated with SB202190 (20 μM) for 6 h to assess the levels of LAMP1, MCOLN1, and TPP1 expression by qRT-PCR. Data represent mean ± SD; \**p*<0.05, \*\**p*<0.01, and \*\*\**p*<0.001.

**Appendix Figure S4. PERK activation by SB202190 reduces the aggregation of APP and α-syn accumulation through TFEB-ALP activation in SH-SY5Y cells related to Figure 6**.

(A) SH-SY5Y cells were transfected with pCAX-APP-Swe/Ind (APP^swe/ind^) for 48 h. Cells were subsequently pretreated with MitoTEMPO (100 nM) for 1 h and then treated with SB202190 (20 μM) for 12 h. Representative image of FL-APP was analyzed by confocal microscopy (*left*) and quantification of FL-APP intensity (*right*). Data represent mean ± SD; \*\*\**p*<0.001. (B, C) SH-SY5Y cells were transiently transfected with α-Syn-A53T for 48 h. Cells were treated with GSK2606414 (1 μM) for 1 h before SB202190 (20 μM) treatment for 12 h. Representative image of α-Syn was detected by confocal microscopy (*left*) and quantification of α-Syn intensity (*right*). Data represent mean ± SD; \*\*\**p*<0.001. (C) The levels of α-Syn and LC3B-II conversion were analyzed by western blotting.

